# Synonymous codon usage regulates translation initiation

**DOI:** 10.1101/2022.05.13.491887

**Authors:** Chloe L. Barrington, Amanda L. Koch, Gabriel Galindo, Emma Larkin-Gero, Evan J. Morrison, Samantha Tisa, Timothy J. Stasevich, Olivia S. Rissland

## Abstract

Synonymous codon usage regulates gene expression such that transcripts rich in optimal codons produce significantly more protein than their nonoptimal counterparts. A major unresolved issue has been understanding the mechanisms by which synonymous codons regulate gene expression. We and others have previously shown that nonoptimal codons slow translation elongation speeds and thereby trigger mRNA degradation. However, differences in transcript abundance are not always sufficient to explain differences in protein levels, suggesting there are additional mechanisms by which codon usage regulates gene expression. Using reporter assays in human and *Drosophila* cells, we found that transcript levels account for less than half of the variation in protein abundance. We demonstrate that the differences at the protein level are not attributable to either protein folding or stability. Instead, we find that mRNAs with nonoptimal codons are bound by fewer ribosomes and that nonoptimal codon usage represses translation initiation. Nonoptimal transcripts are also less bound by the key translation initiation factors eIF4E and eIF4G, providing a mechanistic explanation for their reduced initiation rates. Our results reveal a new mechanism of regulation by codon usage, where nonoptimal codons repress further rounds of translation.

## INTRODUCTION

The open reading frame (ORF) does more than encode for protein: it also regulates gene expression. Our current understanding of the ORF as a major regulatory element represents a shift from the classical view that 5’ and 3’ untranslated regions (UTRs) are the main post-transcriptional regulators of gene expression. Through structures and sequences bound by *trans* regulatory factors, the UTRs can alter the translation, stability, and localization of messenger RNAs (mRNAs) (Grimson et al., 2007, Hinnebusch et al., 2016, Rissland et al., 2017, Leppek et al., 2018, Mayr, 2019). However, we now know that sequences within the ORF also control many of these same processes.

Codon usage is a key way that the ORF regulates gene expression. The genetic code is redundant, meaning that multiple synonymous codons can code for the same amino acid. However, synonymous codons are not used equally. Within each species, some codons are more frequently used, especially in highly expressed genes, and thought to be “optimal.” Other codons are less widely used and are called “nonoptimal” (Grantham et al., 1980, Ikemura, 1985). Synonymous codon usage is an essential and evolutionarily conserved mechanism of gene regulation: a gene with optimal codons produces dramatically more protein than its nonoptimal counterpart, even when the same primary protein sequence is encoded (dos Reis et al., 2004, Liu et al., 2021, Bae and Coller, 2022). This phenomenon underpins the common practice of codon optimization within heterologous expression systems to increase expression of genes of interest (Ikemura, 1981, Gustafsson et al., 2004). Despite the well-known importance of codon optimality on expression levels, understanding the mechanisms by which codon usage impacts gene expression remains an active area of study.

Codon usage can influence nearly every step of the gene expression pathway, but most of its impact occurs through translation. Due to a variety of features, including tRNA abundance and charging, codon usage in the transcriptome, and wobble position interactions, optimal codons are decoded more quickly than nonoptimal ones (dos Reis et al., 2004, Gardin et al., 2014, Gamble et al., 2016, Hanson et al., 2018). Because decoding is often the rate-limiting step in translation elongation (Gardin et al., 2014, Dana and Tuller, 2014), slow decoding slows overall translation elongation. In some species (depending on the nature of the optimal codons), codon usage often changes the GC-content of the transcript, which can also create secondary structural elements that have the potential to curtail ribosome progression (Pop et al., 2014, Mauger et al., 2019, Andrzejewska et al., 2020).

Slow translation elongation speed then affects other aspects of post-transcriptional regulation. One proposed mechanism is that translation elongation rates can alter protein folding (Pechmann and Frydman, 2013, Yu et al., 2015, Komar et al., 1999, Chaney et al., 2017, Fu et al., 2018, Zhou et al., 2015). The evidence underlying this model is that codons encoding linker regions between protein domains are frequently nonoptimal. These are thought to slow ribosomes down between structured elements and allow enough time for proper co-translational peptide folding (Pechmann and Frydman, 2013, Yu et al., 2015, Zhou et al., 2015, Chaney et al., 2017, Liu et al., 2020). Indeed, for this reason, complete optimization can reduce protein yield (Zhou et al., 2013, Jacobson and Clark, 2016, Buhr et al., 2016). Another mechanism by which codon usage can affect gene expression is mediated within the so-called ‘translational ramp,’ which refers to the beginning of the ORF (Tuller et al., 2010, Verma et al., 2019). This region is often disproportionately nonoptimal, which is thought to slow the ribosome, sterically block the start codon, and limit overall initiation rates (Tuller et al., 2010, Chu et al., 2014). These observations have led to the model that the translational ramp acts to space out ribosomes on the ORF (Tuller et al., 2010). Another mechanism by which elongation can regulate gene expression is through surveillance mechanisms. For instance, stalled ribosomes can trigger the ribosome quality control (RQC) pathway, resulting in destruction of the nascent peptide, rescue of the stalled ribosomes, and degradation of the translated mRNA (Brandman, O., and Hegde, 2016, Hickey et al., 2020, Juszkiewicz et al., 2020, Sinha et al., 2020, Wu et al., 2020).

Finally, recent work has shown that slow elongation, specifically through nonoptimal codon usage and slow decoding, triggers mRNA decay by a mechanism distinct from the RQC pathway (Buschauer et al., 2020, Veltri et al., 2021, Mishima et al., 2022). Best described in yeast, this pathway is mediated by the CCR4-NOT deadenylase complex (Presnyak et al., 2015, Hanson et al., 2018, Buschauer et al., 2020). Here, a hallmark of slow decoding (i.e., a ribosome with an empty A and E site) leads to binding of Not5p within the empty E site (Buschauer et al., 2020, Allen et al., 2021). Because Not5p is a component of the CCR4-NOT deadenylase complex, its binding stimulates deadenylation of the mRNA, accelerating the eventual mRNA destruction by the Dhh1p decapping activator, the decapping enzyme, and the 5’è3’ Xrn1 exonuclease (Presnyak et al., 2015, Radhakrishnan et al., 2016, Webster et al., 2018, Buschauer et al., 2020). The connection between synonymous codon usage and transcript stability has now been expanded to a wide array of prokaryotic and eukaryotic organisms, including *E. coli*, *S. pombe*, *Neurospora, Xenopus*, zebrafish, *Drosophila*, mice, and humans, indicating that this mechanism is deeply conserved (Boël et al., 2016, Harigaya and Parker, 2016, Zhou et al., 2013, Zhou et al., 2016, Burow et al., 2018, Bazzini et al., 2016, Mishima and Tomari, 2016, Lampson et al., 2013, Wu et al., 2019, Hia et al., 2019, Narula et al., 2019, Forrest et al., 2020).

Despite the substantial amount of research into how codon usage impacts protein expression, none of these mechanisms can fully explain the repressive effect from nonoptimal codons. This gap in knowledge is most clearly seen in studies on Ras paralogues. Ras genes (HRas, NRas, and KRas) differ in their codon usage, and KRas, the least optimal paralog, produces substantially less protein than HRas, the most optimal one (Lampson et al., 2013, Fu et al., 2018). Importantly, in studies where KRas has been optimized, more protein was produced than could be explained by changes in overall mRNA levels (Lampson et al., 2013, Fu et al., 2018). Similarly, several studies have shown that more ribosomes are bound to optimal mRNAs than nonoptimal ones (Ingolia et al., 2009, Bazzini et al., 2016, Heyer and Moore, 2016, Lima et al., 2017, Fu et al., 2018)—however, this effect is counter to the expectation that, like slow traffic on a highway, slow elongation should result in *more*, not *fewer*, ribosomes on nonoptimal mRNAs. These results raise the possibility that codon usage may affect other post-transcriptional regulatory pathways, including translation itself.

Here, we explored the mechanisms by which codon usage regulates gene expression. Using a series of reporters with large N-terminal tags (to exclude potential effects of the translational ramp), we found that optimal genes produce more protein than nonoptimal ones, beyond what can be explained by changes in mRNA levels. The effect at the protein level can be as large or, in some cases, substantially greater than that at the mRNA level, revealing a robust yet undescribed mechanism by which codon usage regulates gene expression. These effects are conserved from *Drosophila* to humans. The reduction in protein levels cannot be explained by differences in protein stability or folding, nor by premature ribosome termination. Instead, our results indicate that nonoptimal codons repress translation initiation. Moreover, nonoptimal transcripts have reduced binding to key translation initiation factors, eIF4E and eIF4G, suggesting a mechanistic model for how slow elongation represses translation initiation. Together, these results reveal a potent mechanism by which synonymous codon usage regulates gene expression and provide a unifying explanation for many of the longstanding mysteries in the field.

## RESULTS

### Codon optimality affects protein levels beyond what can be explained by mRNA changes

To determine how codon usage affects mRNA and protein levels, we made reporters where an optimal or nonoptimal firefly luciferase coding region was preceded by a large (∼1 kb) N-terminal Flag “spaghetti monster” tag (see below; Viswanathan et al., 2015, Koch et al., 2020). This large tag ensured that any regulation by codon optimality could not be explained by differences in the translational ramp (Tuller et al., 2010). After transfecting these constructs in HEK293T cells along with an optimized *Renilla* luciferase control, we measured transcript levels by RT-qPCR and protein abundance by luciferase assays. As expected, based on recent findings (Wu et al., 2019, Hia et al., 2019, Narula et al., 2019, Forrest et al., 2020), the optimal reporter mRNA was more abundant than its nonoptimal counterpart (2.3 ± 0.7-fold, *p* < 10^-5^; Figure 1A left panel).

**Figure 1.**
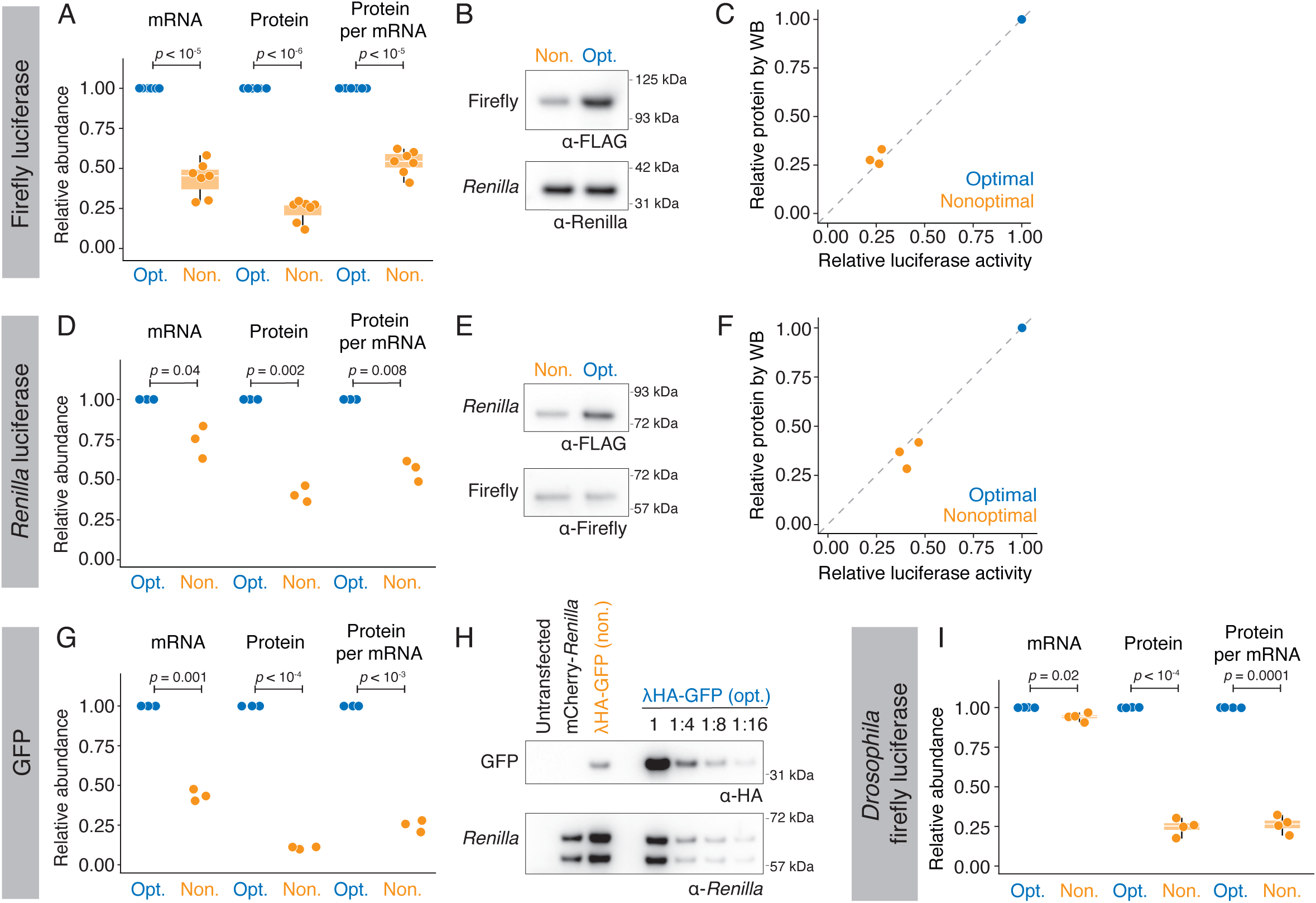
Codon optimality affects protein levels beyond what can be explained by changes at the mRNA level. (A) Firefly luciferase optimality reporters show differences in protein output, even after controlling for mRNA abundance. HEK293T cells were transfected with either Flag spaghetti-monster tagged optimal (“Opt.”) or nonoptimal (“Non.”) firefly luciferase reporters and *Renilla* luciferase, as a control. mRNA abundance was quantified by RT-qPCR, and protein activity was quantified by dual luciferase assay. Shown are box-and-whisker plots for the fold-change in the nonoptimal reporter relative to the optimal one for mRNA abundance, protein abundance, and normalized protein-per-mRNA abundance. P-values were determined using a paired Student’s t-test. (B) Nonoptimal firefly luciferase reporters have reduced protein levels. Western blotting was performed by probing for Flag (to quantify firefly luciferase levels) or for *Renilla* luciferase. (C) Western blotting and luciferase assays show a reduction in firefly luciferase levels due to nonoptimal codons. Shown is a scatter plot comparing the fold-change in the nonoptimal reporter firefly luciferase abundance as determined by luciferase activity or western blotting. There was no significant difference between these measurements, *p* = 0.3 (paired Student’s t-test). (D) As in A, except for spaghetti-monster-tagged *Renilla* luciferase. (E) As in B, except for *Renilla* luciferase. (F) As in C, except for *Renilla* luciferase. *p* = 0.2 (paired Student’s t-test). (G) As in A, except for λHA-tagged GFP, where protein levels were quantified by western blotting. mCherry-*Renilla* was used as a co-transfection control. (H) Nonoptimal GFP shows reduced protein levels. Shown is a representative western blot used to quantify the reduction in λHA-GFP protein levels due to nonoptimal codons. Lysates were serially diluted 1:2 to allow accurate quantitation, and fold-changes were calculated relative to the *Renilla* luciferase loading control. (I) Nonoptimal codons repress protein output in *Drosophila*. S2 cells were transfected with either nonoptimal or optimal λHA-tagged firefly luciferase and *Renilla* luciferase as a co-transfection control. As in A, mRNA and protein levels were determined by RT-qPCR and luciferase assays. P-values were determined using a paired Student’s t-test.

However, differences in protein levels exceeded those at the mRNA level. The optimal protein was 4.2-fold more abundant than the nonoptimal one (*p* < 10^-6^; Figure 1A middle panel). To confirm that these differences could not be explained by protein folding and a corresponding impact on activity, which has been reported to be affected by codon usage (Pechmann and Frydman, 2013, Yu et al., 2015, Zhou et al., 2015, Chaney et al., 2017, Liu et al., 2020), we repeated measurements of protein abundance using western blotting and saw similar results (Figure 1B, C). Together, these data show that there is approximately a two-fold difference in protein expression beyond that at the transcript level (*p* < 10^-5^; Figure 1A right panel). These results were not cell-type specific, as the effect was recapitulated in U2OS cells (Figure S1A). They also did not depend on the addition of optimal *Renilla* luciferase for normalization because results were unchanged when these experiments were repeated with a nonoptimal *Renilla* luciferase (Figure S1B).

To test the generality of this observation, we next asked whether optimization of different reporters gave similar results. We first tested *Renilla* luciferase. As with firefly luciferase, the optimal *Renilla* luciferase mRNA was more abundant than the nonoptimal one (1.37 ± 0.19-fold difference, *p* = 0.04; Figure 1D left panel), but it was even more abundant at the protein level (2.44 ± 0.29-fold difference, *p* = 0.002; Figure 1D middle panel). Activity, as measured by luciferase assay, was similar to total protein abundance, as measured by western blotting (Figure 1E, F). These results thus showed nearly a 2-fold difference in *Renilla* luciferase expression that was unexplained by differences in transcript abundance (Figure 1D right panel). We repeated the experiment using λHA-GFP optimality reporters; these reporters contained a different N-terminal tag, which, while shorter than the original spaghetti monster tag, was still sufficiently long to exclude a possible translational ramp effect (Tuller et al., 2010). Again, the ∼2.3-fold difference in mRNA levels was not sufficient to account for the ∼9.4-fold difference in protein levels, thus revealing nearly an additional four-fold effect at the protein level (*p* < 10^-3^; Figure 1G, H). Thus, our results indicate that nonoptimal codons exert repressive effects beyond changes at the mRNA level in human cells.

We next asked if this effect by poor codon usage was conserved. To do so, we performed paired assays in the Drosophila S2 cell line using λHA-firefly luciferase reporters. Interestingly, although codon optimization had a negligible effect on mRNA abundance (1.06 ± 0.03-fold change, *p =* 0.02; Figure 1I), the optimal reporter was still expressed at nearly four-fold higher abundance at the protein level than the nonoptimal one (*p* = 0.0001; Figure 1I). Thus, four different sets of reporters in both vertebrates and invertebrates showed that poor codon usage affects protein expression more than can be accounted for by changes at the mRNA level. Depending on the reporter and system, the effect at the protein level can be even larger than that seen at the mRNA level.

### Poor codon optimality does not affect protein stability

The observation that codon usage affects protein abundance beyond that which can be explained by mRNA abundance could be due to changes in protein stability or translation. Given that the rate of translation elongation has been proposed to change the dynamics of co-translational folding, and that incorrect folding can target peptides for degradation (Zhou et al., 2013, Jacobson and Clark, 2016, Buhr et al., 2016), we asked if each nonoptimal reporter was less stable than their optimal counterpart by measuring protein half-lives using cycloheximide translation shut-off assays. We first confirmed that cycloheximide effectively inhibited translation with a puromycin incorporation assay (Figure 2A). Nonoptimal and optimal firefly luciferase reporters had a half-life of 15 ± 6 hours and 12 ± 3 hours, respectively; thus, if anything, the nonoptimal reporter was slightly more stable than the optimal one, although these half-lives were not significantly different (*p* = 0.35; Figure 2B). Similar results were obtained with the *Renilla* luciferase reporters, where again the nonoptimal variant had a longer, though not significantly different, half-life than its optimized counterpart (32 ± 17 hours and 18 ± 6 hours, respectively, *p =* 0.18; Figure 2C). As expected, there were no difference in the half-lives of the co-transfection controls (Figure S2A, B).

**Figure 2.**
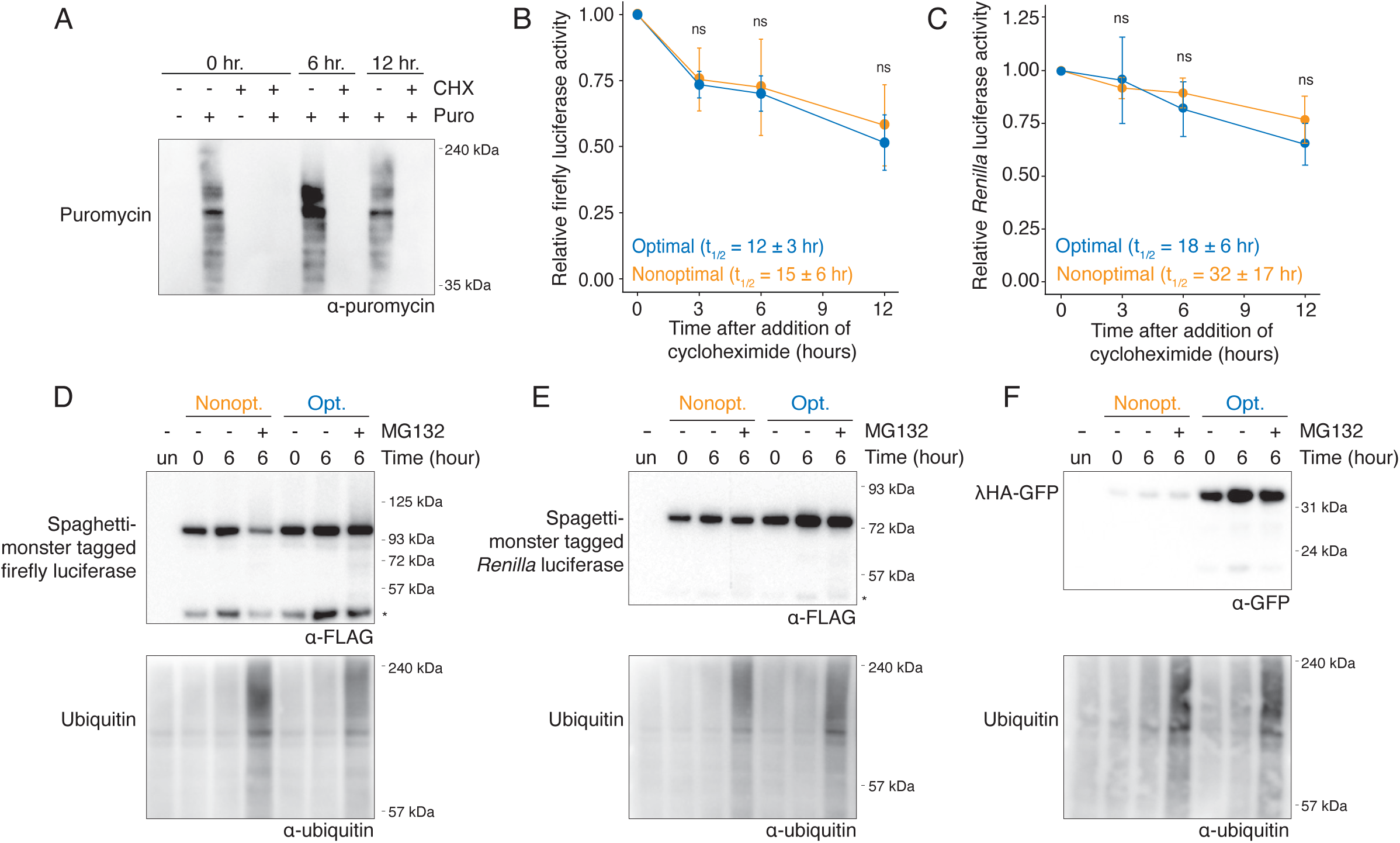
Poor codon optimality does not affect protein stability. (A) Cycloheximide treatment reduces translation. HEK293T cells were treated with cycloheximide or DMSO for zero, six, and 12 hours, and ongoing translation was determined by puromycin incorporation. Puromycin was added ten minutes prior to harvesting, and incorporation was determined by western blotting. (B) Codon optimality does not affect the stability of firefly luciferase. Cells were transfected with the spaghetti-monster firefly luciferase reporters and harvested at various time points after cycloheximide addition. The amount of firefly luciferase was determined by luciferase assayand normalized to the initial (0 hr.) time point. P-values were determined using paired Student’s t-tests. (C) As in B, except for spaghetti-monster *Renilla* luciferase reporters. (D) Codon optimality does not affect proteasome-mediated degradation of firefly luciferase. After cells were transfected with the spaghetti-monster firefly luciferase reporters, they were treated with the proteasomal inhibitor MG132 and harvested at various time points. Western blots were performed, probing for the firefly luciferase reporters (with Flag) and ubiquitin. *, cleavage product (E) As in D, except for spaghetti-monster *Renilla* luciferase. (F) As in D, except for λHA-GFP.

As an alternative approach, we treated the cells with the proteasome inhibitor MG132 to determine whether protein products were ubiquitinated and targeted for degradation. The stabilization of ubiquitinated proteins confirmed that MG132 treatment effectively blocked proteasome activity (Figure 2D–F). However, there was no detectable stabilization of the full-length for either the optimal or nonoptimal luciferase reporters (Figures 2D, E), a result consistent with the relatively long stabilities of these reporters. There was also no stabilization seen from the λHA-GFP reporters (Figure 2F). Taken together, these data indicate that the differences in protein levels caused by codon usage cannot be explained by differences in protein stability.

### Nonoptimal elongation causes association with fewer ribosomes

Because differences in protein stability could not explain how nonoptimal codons affected protein abundance, we turned to translational effects. We first measured the impact of codon usage on translation in bulk using polysome gradients. To minimize technical variation between the reporters, we co-transfected U2OS cells with the spaghetti-monster-tagged firefly luciferase nonoptimal and optimal reporters, and separated polysomes on a 10-60% sucrose gradient (Figure 3A). qRT-PCR (using primers specific to each codon variant) was used to quantify reporter abundance in each fraction (Figure 3B). Compared with the optimal reporter mRNA, a significantly higher proportion of nonoptimal reporter mRNA was present in the monosome fraction (1.38 ± 0.08-fold, *p* = 0.01; Figure 3B, C) and a lower proportion was associated with six or more ribosomes, although this difference was not significant (0.89 ± 0.06-fold, *p =* 0.09; Figure S4A). Thus, the nonoptimal transcript is associated with fewer ribosomes than the optimal one. Although consistent with many other results in the field (Ingolia et al., 2009, Bazzini et al., 2016, Heyer and Moore, 2016, Lima et al., 2017, Fu et al., 2018), these results are contrary to the expectation that because nonoptimal codons slow elongating ribosomes (dos Reis et al., 2004, Gardin et al., 2014, Dana and Tuller, 2014, Gamble et al., 2016, Hanson et al., 2018), there should be more ribosomes on our nonoptimal reporter, not fewer.

**Figure 3.**
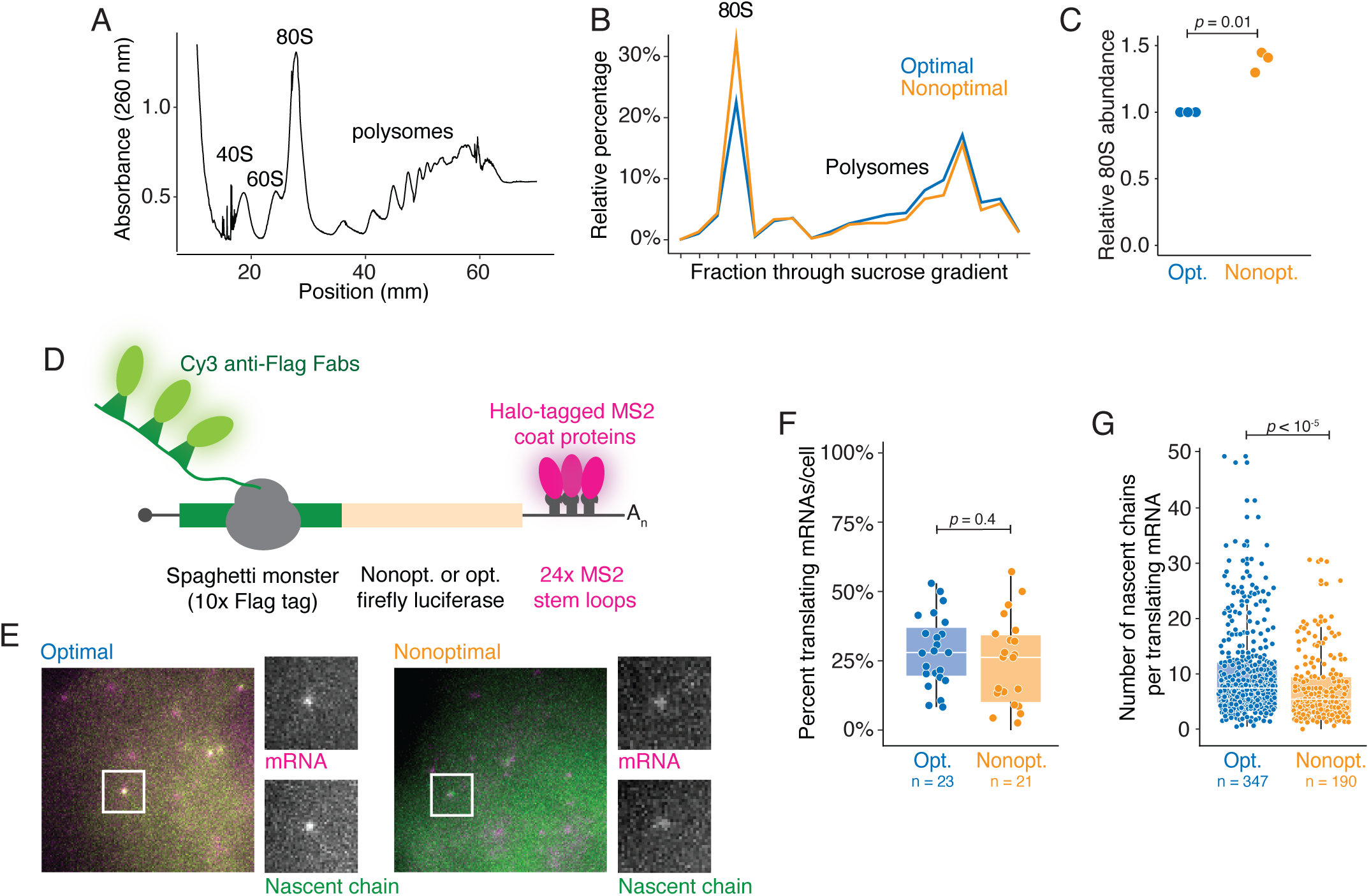
Nonoptimal codons reduces ribosome association. (A) Representative polysome fractionation. HEK293T cells were transfected with both the nonoptimal and optimal firefly luciferase reporters, and lysates were fractionated by sucrose fractionation. Shown is the A260 from a representative gradient. (B) Nonoptimal codon usage affects ribosome association. Fractions from each gradient were pooled based on the number of ribosomes, and the amount of each reporter was quantified with RT-qPCR. Plotted is the percentage of each reporter in each fraction for a representative biological replicate. (C) Nonoptimal codon usage increases association with the 80S fraction. Plotted is the abundance of the nonoptimal firefly reporter in the 80S fraction relative to the optimal reporter. The p-value was determined using a paired Student’s t-test. (D) Schematic of the nascent chain tracking system. Translation is visualized with anti-Flag Fabs conjugated to Cy3, which bind 10x Flag peptide epitopes in the nascent peptide after exit from the ribosome exit tunnel. The mRNA is visualized with Halo-labeled MS2 coat proteins, which bind to 24x MS2 stem loops in the 3’UTR. (E) mRNA and protein signals colocalize for the firefly luciferase reporters. Fluorescent images of representative cells expressing optimal (left) and nonoptimal (right) firefly luciferase reporters. The white boxes highlight single punctum with grey-scale insets, showing both mRNA (MS2-coat proteins) and nascent chain anti-Flag Fab signals. (F) Nonoptimal codons do not affect the likelihood of translation. Plotted is a box-and-whisker plot for the percent of mRNAs translating per cell. Each point refers to a single cell. The p-value was determined using a Mann-Whitney test. (G) Nonoptimal codons reduce the number of ribosomes bound to each mRNA. Plotted is a box-and-whisker plot for the number of nascent chains associated with each translating reporter mRNA. Each point refers to a single mRNA molecule. The p-value was determined using a Mann-Whitney test.

To explore this finding with more resolution, we turned to an *in vivo* imaging system called nascent chain tracking, where active translation on single transcripts can be visualized in living cells (Morisaki et al., 2016, Wu et al., 2016, Wang et al., 2016, Yan et al., 2016, Pichon et al., 2016). In this system, individual reporter transcripts are visualized through 24x MS2 stem loops in their 3’UTR, which are bound by Halo-tagged MS2 coat proteins (Figure 3D). At the same time, a large N-terminal spaghetti monster contains 10x Flag epitopes that are bound co-translationally by fluorescently conjugated anti-Flag antibodies. These reporters are the same as have been used throughout this study.

We first confirmed that translation of both firefly luciferase reporters could be visualized in cells. Plasmid DNA containing each optimality reporter was bead-loaded into U2OS cells, along with purified anti-Flag Cy3-Fab and HaloTag-MCP probes. As expected, for both reporters, we could detect untranslated mRNAs (labeled with the HaloTag alone) and mRNAs undergoing active translation (co-labeled with both HaloTag and anti-Flag Cy3 signals; Figure 3E). To confirm that co-labeled spots were sites of active translation, cells were treated with puromycin, and the anti-Flag Cy3 signal rapidly dissociated from each mRNA punctum (Supplemental Movies 1 & 2). In other words, this system can detect translation of the firefly luciferase reporters in living cells with single-molecule resolution.

With the ability to observe active translation on single mRNA molecules, we next examined the impact of codon optimality on translation (Supplemental Movies 3 & 4). Given that nonoptimal codon usage leads to reduced protein output and decreased association with polysomes by bulk analysis, we envisioned two, non-mutually exclusive possibilities. The first is that nonoptimal codon usage might prevent translation altogether—that is, reduce the likelihood of translation occurring on a given mRNA. To investigate this hypothesis, we measured how many individual reporter mRNAs in each cell, at steady state, co-localized with nascent chain signal. Based on this co-localization, we observed no difference in the fraction of mRNAs translated per cell (optimal reporter: 28.5 ± 12.9%; nonoptimal reporter: 23.8 ± 16.3%; *p* = 0.4; Figure 3F). These data suggest that codon usage does not change the probability that an mRNA will be translated.

The second possibility is that the magnitude of translation itself is dampened, thus reducing the number of initiating ribosomes per translation burst. To explore this second possibility, we measured the intensity of anti-Flag signal associated with each translating reporter (Figure 3G), This intensity score corresponds to the number of nascent peptides actively synthesized on each mRNA molecule, and intensity values were normalized using Flag-tagged β-actin constructs (Koch et al., 2020). There was a significant difference in intensity (and thus the number of ribosomes translating) between the two reporters (*p* < 10^-5^; Figure 3G). On the optimal reporter, the intensity corresponded to a median of 7.4 nascent chains, while the same measurements on the nonoptimal reporter showed a median of 5.4 nascent chains. Therefore, single-molecule imaging data supports our previous polysome profiling results: nonoptimal codon usage reduces the number of translating ribosomes.

### Poor codon usage does not cause incomplete translation

One possibility that could explain the reduced number of translating ribosomes on nonoptimal transcripts is that poor codon usage leads to ribosome drop-off. One way this could occur is through frameshifting, which has been proposed to take place on stretches of nonoptimal codons (Celik et al., 2017). It could also occur through a mechanism analogous to what happens no-go decay (NGD) and RQC, where coding regions stall the ribosome and ultimately result in ribosome splitting and destruction of the nascent peptide. In theory, nonoptimal codons could act as a RQC substrate and slow ribosome progression enough to cause stalls and trigger surveillance mechanisms. Two major signatures of these mechanisms would be 1) proteasomal degradation of truncated nascent peptides and 2) ribosome drop-off.

We first tested the extent to which nonoptimal codon usage leads to production of truncated protein products. Because these products are likely unstable, we transfected cells with either optimal or nonoptimal versions of various reporters (spaghetti-monster tagged firefly luciferase, spaghetti-monster tagged *Renilla* luciferase, and λHA-GFP) and then treated the cells with MG132, as before. However, even when western blots were overexposed, there was no evidence of small fragments that accumulated specifically with any of the nonoptimal reporters (Figures 4A–C). These data suggest that truncated protein products are not produced in response to nonoptimal codon usage.

**Figure 4.**
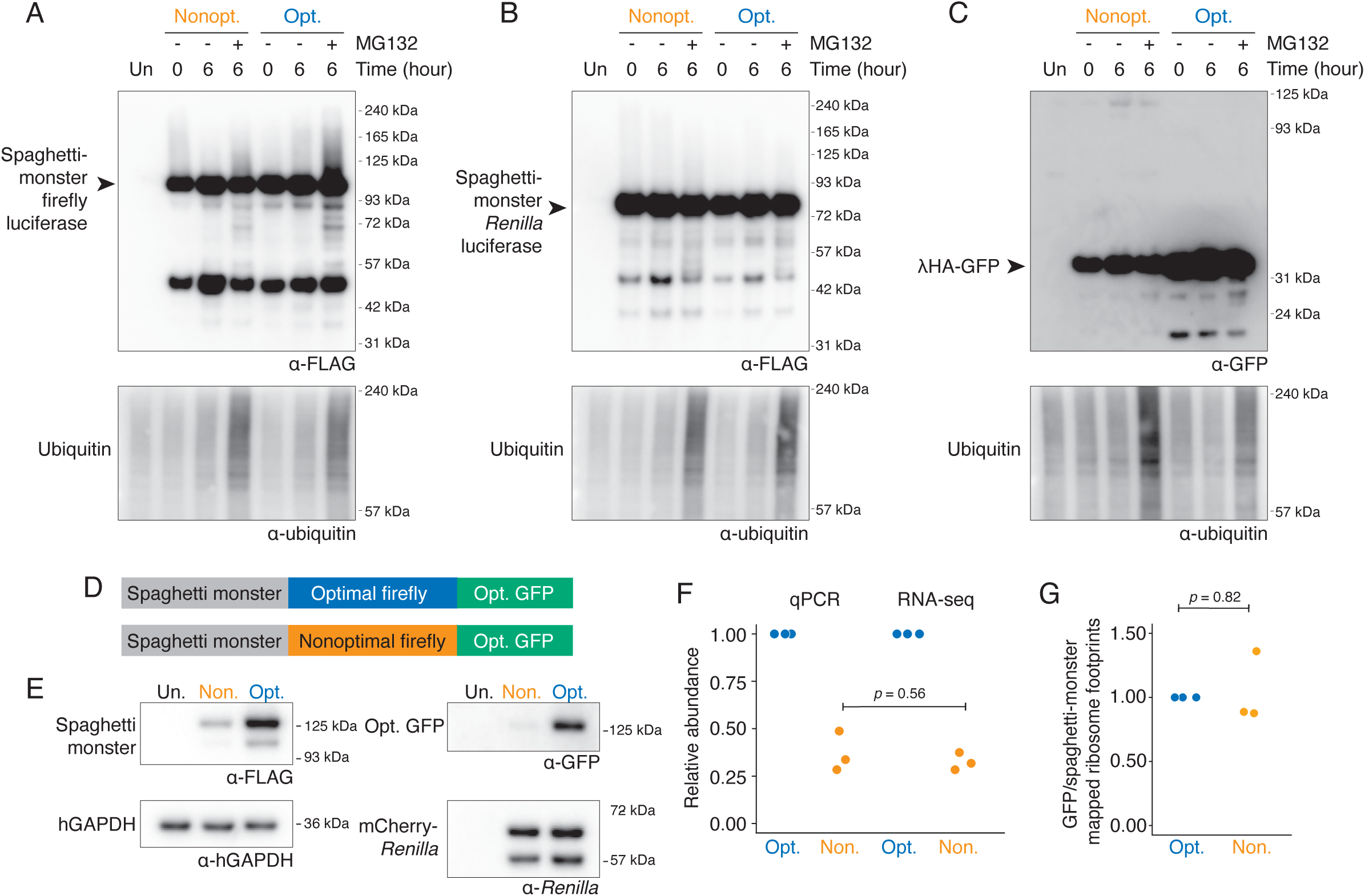
Nonoptimal codons do not lead to incomplete translation. (A) Nonoptimal codons do not lead to truncated protein products. HEK239T cells were transfected with the spaghetti-monster firefly luciferase reporters and treated for six hours with MG132 or DMSO. Western blotting was performed, probing for the firefly luciferase reporter (with α-Flag) and for ubiquitin. Twice the volume of lysate was loaded for samples with nonoptimal firefly luciferase. Shown is an overexposed blot to detect the accumulation of truncated protein products. (B) As in A, except for the spaghetti-monster *Renilla* luciferase reporters. Twice the volume of lysate was loaded for samples with nonoptimal *Renilla* luciferase. (C) As in A, except for the λHA-GFP reporters. Four times the volume of lysate was loaded for samples with nonoptimal λHA-GFP. (D) Schematic of the dual-tagged reporters used for the Ribo-seq experiment. (E) Nonoptimal codons repress protein accumulation for the dual-tagged firefly luciferase reporters. HEK293T cells were transfected with the dual-tagged reporters and mCherry-*Renilla* luciferase, as a co-transfection control. Protein abundance was determined by western blotting, probing for the N-terminal spaghetti monster tag (with α-Flag), the C-terminal tag (with α-GFP), *Renilla* luciferase, and hGAPDH. (F) Nonoptimal codons reduce the mRNA abundance of the dual-tagged reporters. mRNA abundance for each reporter was determined by RT-qPCR or RNA-seq (from the same samples used for the Ribo-seq libraries) and normalized to the *Renilla* luciferase co-transfection control. The p-value was determined by an unpaired Student’s t-test. (G) Nonoptimal codons do not lead to ribosome drop-off. Shown is the relative ratio of reads mapping to the N-terminal spaghetti monster tag and the C-terminal GFP tag for the optimal and nonoptimal reporters. The first 25 codons of the spaghetti-monster tag and the last 25 codons of the GFP tag were excluded from the analysis. The p-value was determined using a paired Student’s t-test.

We next wanted to look directly at ribosome drop-off using ribosome profiling. To do so, we modified our spaghetti-monster tagged firefly luciferase reporter such that it was now followed by a C-terminal optimized GFP (Figure 4D). Importantly, both reporters are identical except for the synonymous codons used in the internal firefly luciferase region. Consistent with the previous single-tagged reporters (Figure 1), nonoptimal codon usage reduced mRNA abundance (2.7-fold, *p =* 0.009) and reduced protein expression even further (5.8-fold, *p =* 0.0004), as determined by both western blotting and dual luciferase assays (Figure 4E, Figure S3A). Thus, the dual-tagged reporters recapitulate the biology observed earlier with our single-tagged reporters.

We performed RNA- and Ribo-seq on cells transfected with these reporters in biological triplicate. As expected, RNA-seq measurements showed a decrease in nonoptimal mRNA abundance that was indistinguishable from previous qRT-PCR measurements (*p* = 0.56; Figure 4F, Figure S3B). To directly investigate drop-off, we determined the ratio of ribosome-associated reads mapping to the tags that preceded or followed the firefly luciferase coding region (spaghetti monster and GFP, respectively). However, there was no difference in this ratio between the two reporters (*p* = 0.82; Figure 4G). Moreover, there was no evidence of substantial frameshifting due to the nonoptimal firefly luciferase coding region (Figure S3C). Hence, premature termination cannot explain the difference in ribosome occupancy between the optimal and nonoptimal reporters.

### Poor codon usage inhibits translation initiation

The other possibility for the reduced number of ribosomes on our nonoptimal reporters is reduced translation initiation. To test this possible mechanism, we referred to our Ribo-seq datasets (Figure 4). Reasoning that a decrease in translation initiation would lead to a decrease in translation *before* the nonoptimal region, we determined the translational efficiency of the spaghetti-monster tag upstream of the firefly luciferase coding region. Here, we defined translation efficiency as the number of Ribo-seq reads normalized to RNA-seq reads. As a control, we also calculated the translational efficiency of the co-transfected *Renilla* luciferase, which did not significantly differ between the conditions (*p =* 0.50; Figure S4B). In contrast to the control reporter, the translational efficiency of the upstream spaghetti-monster tag was significantly reduced in the context of the nonoptimal reporter relative to the optimal one (*p* = 0.02; Figure 5A). Because the ribosome encounters this tag before the nonoptimal coding region, this result strongly supports a model of reduced translation initiation on the nonoptimal reporter.

**Figure 5.**
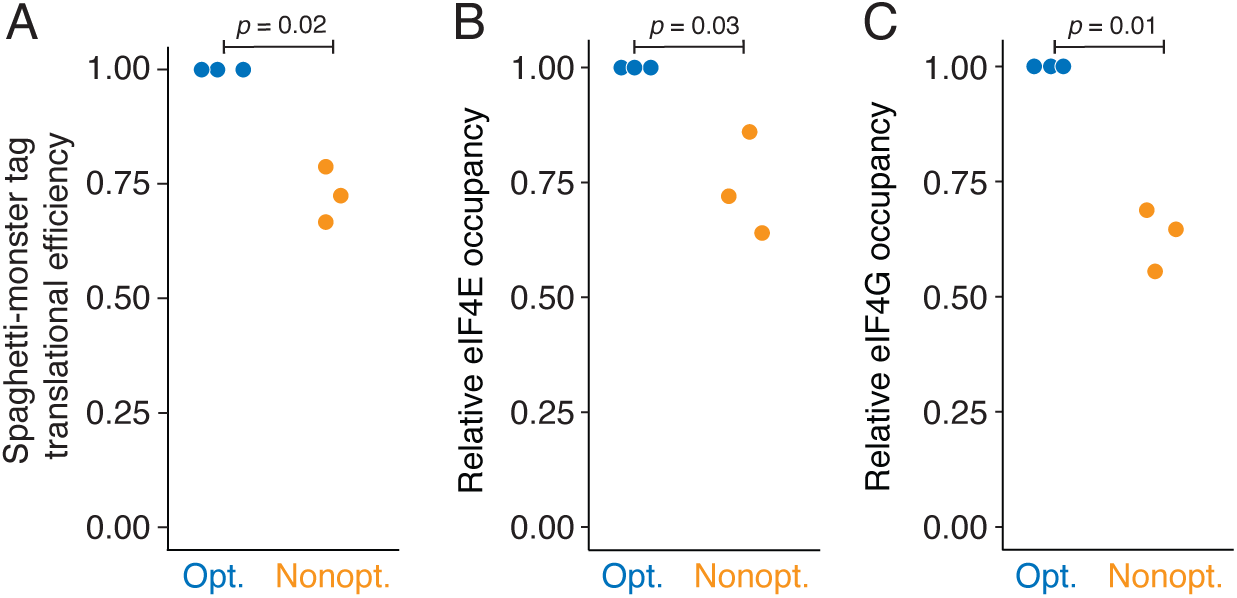
Nonoptimal codons inhibit translation initiation. (A) Nonoptimal codons reduce upstream ribosome association. Shown is the translational efficiency (defined as Ribo-seq reads relative to RNA-seq abundance) in the N-terminal spaghetti-monster tag, which is upstream of the optimal or nonoptimal firefly luciferase coding region, in the dual-tagged reporters. The optimal reporter was normalized to one using the co-transfected *Renilla* luciferase control. The p-value was determined by a paired Student’s t-test. (B) Nonoptimal codons reduce eIF4E binding. HEK293T cells were transfected with the optimal or nonoptimal spaghetti-monster firefly luciferase reporters and a *Renilla* luciferase control. eIF4E RNA immunoprecipitations were performed, and the relative fraction of each reporter pulled down was determined by qRT-PCR. The occupancy on the optimal reporter was normalized to one, using *Renilla* luciferase. The p-value was determined using a paired Student’s t-test. (C) As in B, except for eIF4G RNA immunoprecipitations.

To explore the molecular mechanism underlying the repressed translation initiation, we investigated the binding of two core eukaryotic translation initiation factors: eIF4E, the cap binding protein, and eIF4G, the large scaffold protein. We performed RNA immunoprecipitations with these two proteins in cells transfected with one of the two spaghetti-monster tagged firefly luciferase optimality reporters along with a *Renilla* luciferase transfection control. We confirmed that each protein was pulled down using western blotting (Figure S4C, D), and determined the extent to which each reporter was immunoprecipitated using qRT-PCR. To control for technical differences in transfection and pull-down efficiency, we normalized each firefly luciferase reporter to the *Renilla* luciferase control. Consistent with nonoptimal codons repressing translation initiation, the fraction of immunoprecipitated nonoptimal reporter mRNA was significantly reduced for both eIF4E and eIF4G (*p* = 0.03 and 0.01, respectively; Figure 5B, C). Thus, these data indicate that nonoptimal codon usage reduces protein expression by repressing translation initiation via a molecular mechanism that culminates in reduced eIF4E and eIF4G binding.

## DISCUSSION

It has been known for more than four decades that synonymous codon usage dictates gene expression, and this observation has provided a rationale for codon-optimizing transcripts in heterologous expression systems. However, the mechanisms by which synonymous codons modulate gene expression are still not fully understood. In addition to potentially affecting protein folding and transcription, we and others have shown that by slowing decoding rates of the elongating ribosome, poor codon usage leads to decreased mRNA stability; this mechanism appears to be deeply conserved and has been observed in bacteria and a wide variety of eukaryotic species (Boël et al., 2016, Harigaya and Parker, 2016, Burow et al., 2018, Bazzini et al., 2016, Mishima and Tomari, 2016, Lampson et al., 2013, Wu et al., 2019, Hia et al., 2019, Narula et al., 2019, Forrest et al., 2020). However, there have been hints that changing mRNA abundance may not be sufficient to explain the effects of nonoptimal codons.

Here, we show that nonoptimal codons also repress translation initiation (Figure 6). Using a variety of reporters in two different species, we show that changes at the mRNA level are not sufficient to explain changes at the protein level (Figure 1). Importantly, for all our reporters, we included a large N-terminal tag to exclude the potential impact of nonoptimal codons on the translational ramp, where slow initial elongation could sterically block start codon recognition by trailing ribosomes. We show that the differences in protein levels cannot be explained by differences in protein activity (which could reflect altered folding) or protein stability (Figure 2). Instead, as we show with a combination of bulk and single-molecule assays, nonoptimal codons reduce ribosome association (Figure 3). The reduced ribosome association is not due to increased ribosome drop-off or incomplete translation (Figure 4), but instead is caused by reduced translation initiation (Figure 5). Indeed, the proximal cause of reduced translation initiation appears to be inhibited binding of the translation initiation factors eIF4E and eIF4G. While reduced initiation has been observed in *Neurospora*, but the proposed mechanism of action, phosphorylation of eIF2α (Lyu et al., 2021), cannot explain the mechanism we observe in human cells. Taken together, our results reveal an additional, potent post-transcriptional mechanism by which codon usage regulates gene expression in metazoa.

**Figure 6.**
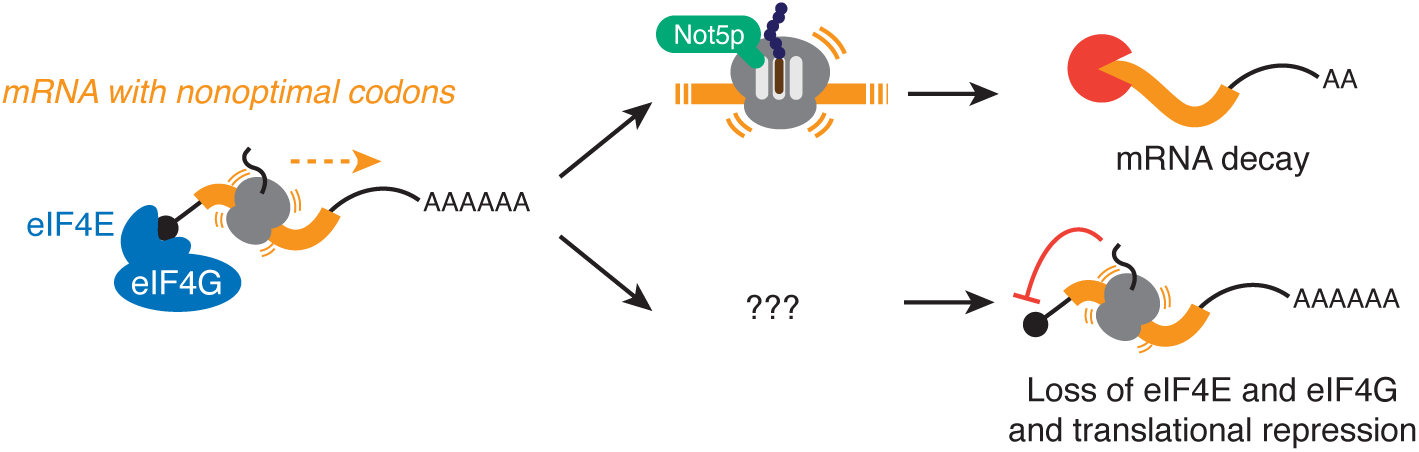
Model: Nonoptimal codons repress gene expression by two post-transcriptional mechanisms. Nonoptimal codons slow translation elongation and can lead to a ribosome state with an empty A- and E-site. In yeast, this state can be sensed by Not5p and thus recruits the CCR4-NOT deadenylase complex to trigger mRNA destabilization. Slow elongation also represses translationinitiation, likely by reducing eIF4E and eIF4G binding. It is unclear what ribosome state is sensed—and how this information is transmitted—to repress translation initiation. The relationship of mRNA destabilization and translation initiation, as caused by nonoptimal codons, is also unknown.

Our results resolve a contradiction in the field. Many studies in a variety of model systems (including yeast, zebrafish, and human cells) have observed that nonoptimal codon usage decreases ribosome association (Ingolia et al., 2009, Bazzini et al., 2016, Heyer and Moore, 2016, Lima et al., 2017, Fu et al., 2018). However, if codon usage only slowed translation elongation, the opposite should be true. This situation is analogous to traffic jams where slow speeds lead to more, not fewer, cars on the highway. However, our finding that nonoptimal codons repress translation initiation reconcile this contradiction. Fewer ribosomes are associated with nonoptimal transcripts, despite the slower elongation speeds, because fewer ribosomes are loaded on the transcripts. In a sense, this result suggests that, in the cell, slow-downs due to nonoptimal codons may lead to a response similar to “on-ramp metering” that motorists experience during rush hour. Thus, our findings reconcile a long-standing discrepancy between slow decoding speeds and reduced ribosome number.

Although we investigate several reporters in our study, one limitation is that we have not formally shown whether nonoptimal codons reduce translational efficiency transcriptome-wide. Of course, to do so on a transcriptome-wide scale is inherently fraught with caveats because of the co-evolution of UTRs and coding regions—genes with poor codon usage often have long UTRs associated with elements that repress translation and so reduce the potential of nonoptimal codons to exert an effect (Giraldez et al., 2006, Mishima and Tomari, 2016, Medina-Muñoz et al., 2021). However, the discrepancy between ribosome association and slow elongation has been noted in endogenous genes, in humans and otherwise (Ingolia et al., 2009, Bazzini et al., 2016, Heyer and Moore, 2016, Lima et al., 2017, Fu et al., 2018), strongly suggesting that this phenomenon is not unique to our constructs. Our study also lacks the resolution to determine what step(s) of initiation are affected, although our results point to eIF4E and eIF4G acting as the proximal cause for reduced translation initiation.

Our findings suggest several new questions (Figure 6). First, what is the mechanism by which nonoptimal codons repress translation initiation? Although formally it is possible that optimal codons could bolster initiation rates, given the link between nonoptimal codons and mRNA instability, we prefer a model whereby slow elongation causes a distinct ribosome state that represses translation. There are two obvious possible “slow” ribosome states that could be sensed: a ribosome with an empty A- and E-site or transiently collided ribosomes. The factors that are reported to bind to these states all have the potential to repress translation initiation. For instance, in yeast, Not5p binds the empty E-site and helps recruit deadenylation machinery; given the link between deadenylation and translational repression, it could be that Not5p (CNOT3, in humans) also represses translation initiation (Buschauer et al., 2020, Veltri et al., 2021, Mishima et al., 2022). An alternative is the recruitment of EDF1/GIGYF2/4EHP, which are known to recognize collided ribosomes as part of the RQC pathway and repress translation initiation on those substrates (Brandman, O., and Hegde, 2016, Hickey et al., 2020, Juszkiewicz et al., 2020, Sinha et al., 2020, Wu et al., 2020). Determining which ribosome state is sensed and how this state is transmitted to the translation machinery is an important next step. Such experiments will also uncover the relationship between mRNA decay and translational repression that is triggered by nonoptimal codons.

Our findings have revealed a new mechanism by which nonoptimal codon usage represses gene expression: reduced translation initiation. This effect is conserved at least from *Drosophila* to humans. Strikingly, this effect is as great, if not greater, than the effect of nonoptimal codons on mRNA levels, and so repressing translation initiation is potent way by which nonoptimal codons regulate gene expression. Because of the recent reliance on mRNA levels to investigate the impact of codon optimality on gene expression, our results explain why this effect has not been recognized until our study. These findings suggest that elongation and initiation machinery tag- team to regulate gene expression and raise important mechanistic questions about their cooperation for future studies.

## ACKNOWLEDGEMENTS

We thank Dr. Suja Jagannathan, Dr. Matt Taliaferro, Dr. David Bentley, and other members of the Rissland lab for thoughtful and helpful discussions. We thank Dr. Neel Mukherjee for sharing several Ribo-seq reagents and help with Ribo-seq data processing. We also thank Amber Baldwin and Kathryn Walters in the Mukherjee lab for their expertise (and patience) when helping with the Ribo-seq protocol and analysis. This work was supported by NIH grant R35GM128680 (OSR) and the RNA Bioscience Initiative, University of Colorado School of Medicine (OSR), NIH training grant T32-GM136444 (CLB), and NSF grant MCB-1845761 (TJS).

## AUTHOR CONTRIBUTIONS

**Chloe Barrington:** Conceptualization, Methodology, Software, Formal analysis, Investigation, Writing – Original Draft, Review & Editing, Visualization, Supervision. **Amanda Koch:** Validation, Formal analysis, Investigation, Writing – Review & Editing. **Gabriel Galindo:** Validation, Formal analysis, Investigation, Writing – Review & Editing. **Emma Larkin-Gero:** Investigation, Writing – Review & Editing. **Evan Morrison:** Investigation, Writing – Review & Editing. **Samantha Tisa:** Investigation, Writing – Review & Editing. **Timothy Stasevich:** Methodology, Resources, Data curation, Writing – Review & Editing, Supervision, Project administration, Funding acquisition. **Olivia Rissland:** Conceptualization, Methodology, Writing – Review & Editing, Visualization, Supervision, Project administration, Funding acquisition. The Authors declare that there is no conflict of interest.

## STAR METHODS

### RESOURCE AVAILABILITY

#### Lead contact

Further information and requests for resources and reagents should be directed to and will be fulfilled by the lead contact, Olivia Rissland.

#### Materials Availability

Plasmids generated in this study will be deposited with Addgene.

#### Data and Code Availability

Ribo-seq and RNA-seq data have been deposited on GEO (GSE202900).

### EXPERIMENTAL MODEL AND SUBJECT DETAILS

#### Cell lines

HEK293T and U2OS cells were grown in DMEM with high glucose and pyruvate (Gibco), supplemented with 10% FBS (Gibco) in a humidified incubator maintained at 37**°**C with 5% CO_2_. Cells were transfected using Lipofectamine 2000 (Invitrogen) and harvested 24 hours after transfection.

S2 cells were maintained in serum-free ExpressFive media (Gibco) supplemented with 20 mM L-glutamine (Gibco) and kept in a humidified incubator at 28**°**C. S2 cells were transfected using Effectene reagent (Qiagen) and harvested 48 hours after transfection.

### METHOD DETAILS

#### qRT-PCR

Cells were washed three times with cold PBS before adding 1 mL TRIzol Reagent (Invitrogen), and RNA was extracted according to manufacturer’s instructions. 1 µg RNA was treated with TURBO DNase (Invitrogen) and reverse transcribed with random hexamer priming (Invitrogen) and Superscript IV Reverse Transcriptase (Invitrogen). qPCR was performed with iTaq Universal SYBR Green Supermix (Bio-Rad) on a LightCycler 480 (Roche) machine. Cycling conditions were as follows: 15 min at 95°C, and then cycled 35x at 95°C for 15 sec, 65°C for 30 sec, and 72°C for 30 seconds. Signal acquisition was performed after each cycle. Samples were measured in technical duplicate and at least three biological replicates were performed per experiment.

#### Dual luciferase assays

Cells were lysed with 1x passive lysis buffer from dual-luciferase reporter assay system kit (Promega), according to the manufacturer’s instructions. 20 µL lysate was added to 96-well flat bottom plate (Greiner Bio-One) in technical triplicate, before performing the dual luciferase assay on a Glomax Navigator plate reader (Promega). Relative luciferase enzymatic activity was quantified by taking the ratio of luciferase values, and the geometric mean was calculated between technical triplicates.

#### Puromycin incorporation assay

Cells were treated with 80 µg/mL cycloheximide for a variable amount of time, before the media was replaced to contain 2 µg/mL puromycin for ten minutes. Puromycin incorporation into nascent peptides is a readout of active translation, while the lack of puromycin incorporation indicates translational shut-off.

#### RNA immunoprecipitation

EZview Red Protein G Affinity Gel beads (Millipore) were prepared by washing twice with lysis buffer A (100 mM KCl, 0.1 mM EDTA, 20 mM HEPES-KOH, pH 7.6, 0.4% NP-40, and 10% glycerol, along with freshly added SUPERase•In [Ambion], 1 mM DTT, and complete mini EDTA-free protease inhibitor cocktail tablets [Roche]). Beads were resuspended in lysis buffer A and blocked with salmon-sperm DNA (Sigma), rotating overnight at 4**°**C. The next day, cells were washed three times with cold PBS pH 7.2 (Gibco) and lysed in cold lysis buffer A, clarified by spinning 21,000 x g at 4**°**C for five minutes, and 50 µL was removed as input. The remaining lysate was incubated with the appropriate antibody for one hour, rotating at 4**°**C. Blocked EZview Protein G beads were washed three times with lysis buffer A before the slurry was added to each IP. Lysates were further incubated for one hour, rotating at 4**°**C. The beads were washed three times with lysis buffer A, transferring to new tubes after the first wash, and split for future analysis: TRIzol reagent (Invitrogen) was added to 90% and processed for qRT-PCR, while the remaining 10% were prepared for western blotting to confirm immunoprecipitation efficiency. Both eIF4E and eIF4G antibodies were purchased from MBL International (2.5 µL and 6 µL used per IP, respectfully), and rabbit polyclonal IgG control antibody was purchased from SinoBiological (CR1, 7.5 µL per IP).

#### Western blotting

Lysates were prepared with NuPAGE 4x LDS Sample buffer (Invitrogen) and 10x Bolt Sample Reducing Agent (Invitrogen) and boiled at 95**°**C for five minutes. Samples were run on NuPAGE 4-12%, Bis-Tris, 1.0 mM, Mini Protean precast gels (Invitrogen) with 1x MES SDS Western Running Buffer (Invitrogen). The protein was transferred to a PVDF membrane (Amersham), according to the manufacturer’s instructions. The membrane was blocked with PBST (1x PBS and 0.1% Tween) with 5% nonfat powdered milk and rocked at room temperature for thirty minutes. Antibodies were diluted in PBST with 5% milk, and blots were incubated rocking overnight at 4**°**C. The following day, the membranes were washed four times with PBST and incubated with light-chain specific HRP-linked rabbit and mouse (Abcam) secondary antibodies (except for eIF4E, where HRP-linked rabbit [Cell Signaling] secondary was used). Membranes were washed four times with PBST. Protein signal was detected with ECL (Cytiva) and imaged on Sapphire Western blot imager (Azure). Western intensity was quantified using Fiji ImageJ. Western blotting was performed in biological triplicate.

#### Sucrose gradients

U2OS cells were co-transfected with both spaghetti-monster tagged firefly luciferase optimality reporters. Just prior to harvesting, cycloheximide was added to 100 µg/mL final concentration (Sigma), and cells were incubated at 37°C for ten minutes. All subsequent harvesting steps were done on ice with cold buffers. The cell monolayer was washed twice with PBS + CHX (100 µg/mL), and 500 µL polysome lysis buffer (10 mM Tris-HCl, pH 7.4, 5 mM MgCl2, 100 mM KCl, and 1% Triton X-100, along with freshly added 2 mM DTT, 500 U/mL SUPERase•In [Ambion], 100 µg/mL cycloheximide, and complete mini EDTA-free protease inhibitor cocktail tablets [Roche]), was added per 10-cm plate. Lysed cells were collected by scraping, sheared four times with a 26-gauge needle, and centrifuged at 1300 x g for ten minutes at 4°C to clarify. Lysates were flash-frozen in liquid nitrogen and stored at –80°C.

10% and 60% sucrose gradient solutions were made with gradient buffer base (20 mM HEPES-KOH, pH 7.4, 5 mM MgCl2, and 100 mM KCl) and filter-sterilized using 0.22-micron PES filter (Millipore). Just prior to pouring, 2 mM DTT, 500 U/mL SUPERase•In (Ambion), and 100 µg/mL cycloheximide was added. 10%-60% gradients were made using the Biocomp Gradient Station per pre-programmed instructions into Open-Top Polyclear centrifuge tubes (Seton Scientific) and stored at 4°C for at least an hour. Lysate was loaded onto the gradient and spun at 36,000 rpm for two hours at 4°C with SW41 TI rotor. Sixty fractions were collected per sample using Piston Gradient fractionator with Triax Full Spectrum Flow Cell. Those fractions were pooled based on ribosome density and brought to 750 µL with RNase-free water. RNA was isolated using phenol-chloroform extraction and precipitated at –80°C overnight with ethanol and sodium acetate. The extracted RNA was processed for qRT-PCR. The percentage of both optimality reporter transcripts in each fraction were calculated. Sucrose gradients were performed in biological triplicate.

#### Single-molecule imaging

U2OS cells were plated on 35 mm glass-bottom MatTek dishes at ∼80% confluency in DMEM (Thermo Fisher Scientific) with 10% (v/v) FBS and 1mM L-Glutamine (DMEM+) the day prior to imaging. On the day of imaging, cells were bead loaded with the nascent chain tracking components: 1.5 µg reporter plasmid DNA (either spaghetti-monster-tagged firefly luciferase nonoptimal and optimal reporters, with 24x MS2 stem loops in the 3’UTR), 100 µg/mL Cy3- labeled α-Flag Fab, and 33 µg/L purified HaloTag-MCP protein. Cells were washed three times with phenol red free DMEM and the media was changed with phenol red free DMEM+ one hour after bead-loading to remove beads. Cells were stained with 200 nM JF646-HaloTag ligand in one mL phenol red free DMEM+ 30 minutes prior to imaging. After ligand incubation, cells were washed three times with phenol red free DMEM and the media was replaced with two mL phenol red free DMEM+. Imaging was done 5-7 hours after bead-loading.

Images were taken on a custom-built microscope (Morisaki et al, 2016). Briefly, the microscope is equipped with 488, 561 and 637 nm solid-state lasers (Vortran), all coupled and focused on the back focal plane of the objective (60×, NA 1.49 oil immersion objective, Olympus). Fluorescent emission signals are split using an imaging grade, ultra-flat dichroic mirror (T660lpxr, Chroma) and the emission beams are focused with a 300 mm tube lens onto two aligned EM-CCD cameras (iXon Ultra 888, Andor). The combination of the imaging objective, 300 mm tube lens, and EM-CCD camera sensors produce 100× images with 130 nm/pixel. One camera detects far-red fluorescence (in this case corresponding to mRNA marked by JF646 HaloTag-MCP), while the other detects red fluorescence (in this case corresponding to translation marked by Cy3- conjugated Fab). The far-red signal is excited with the 637 nm laser using a 731/137 nm emission filter (FF01-731/137/25, Semrock), while the red signal is excited with the 561 nm laser using a 593/46 nm emission filter (FF01-593/46-25, Semrock). Signal-to-noise is enhanced by operating the microscopy in HILO (highly inclined and laminated optical sheet) mode (Tokunaga et al., 2008).

The lasers, filter wheel, cameras, and the piezoelectric stage were all synchronized by an Arduino Mega board (Arduino). The exposure time of the cameras was 53.64 msec throughout the experiments. The camera’s read-out time from the combination of imaging size, readout mode, and the vertical shift speed was 23.36 msec, resulting in an imaging rate of 13 Hz (77 msec per image). The excitation laser lines were digitally synched to ensure they only illuminated cells when the camera was exposing to avoid excessive photobleaching. Laser powers for all images were 20 mW for 637 nm and 5 mW for 561 nm with an ND10 neutral density filter at the beam expander.

Before imaging, live cells were placed into an incubation chamber (Okolab) at 37 °C with 5% CO_2_ on a piezoelectric stage (PZU-2150, Applied Scientific Instrumentation). The focus was maintained with the CRISP Autofocus System (CRISP-890, Applied Scientific Instrumentation). Image acquisition was performed using opensource Micro-Manager (Edelstein et al. 2014). Cell volumes (comprising 13 z-stacks with 0.5 µm steps between each stack) were acquired at full speed, with an imaging delay set so that consecutive volume acquisitions were separated by a six second interval. Cells were selected for imaging if they exhibited at least ten translating mRNA. For puromycin experiments, five time points were taken prior to the addition of 0.5 mL of DMEM+ containing puromycin to achieve a final concentration of 50 µg/mL puromycin.

For image analysis, collected images were first pre-processed with Fiji (Schindelin et al., 2012). First, a 2D maximum intensity projection was generated from the original 3D movies, followed by background subtraction (the background was chosen from the dimmest cell- and debris-free regions of images). Post-processed images were then analyzed using a custom Mathematica (Wolfram Research) routine to detect and track particles in the RNA channel (red color). In this code, images are band-pass filtered so the positions of the intensity centroid of single mRNA spots can be easily detected using the built-in Mathematica routine ComponentMeasurements. Particles detected were then linked through time if they were within 5 pixels (650 nm) of one another in consecutive frames. Particle tracks lasting at least five frames were selected for further analysis.

For each frame of each track, crops (15 pixels x 15 pixels) centered on the registered mRNA coordinate were made and averaged through time. Again, using Mathematica’s built-in bandpass filter and ComponentMeasurements command, the time-averaged crops corresponding to each track were categorized based on their signals in the red (mRNA) and green (translation/nascent chains) channels. If particles were detected in the red channel only, the spot was categorized as a ‘non-translating’ spot. If particles were detected in both the red and green channels, the spot was categorized as a ‘translating’ spot. After categorizing spots in this automated fashion, all spots were again hand-checked to minimize error. Finally, the original 2D maximum intensity projected images corresponding to each hand-checked track were fit to a 2D Gaussian to determine their precise XY-coordinates and signal intensities (using the built-in Mathematica routine NonlinearModelFit). The 2D Gaussian used for fitting had the following form:

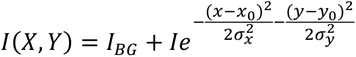

where *I_BG_* is the background signal intensity, *I* the particle peak intensity, (*σ_x_*, *σ_y_*) the spread of the particle signal, and (*x*_0_, *y*_0_) the particle location. From these data, the intensity, position through time, and number of spots over time in each track were quantified.

To correct for any positional offset between the two cameras, Mathematica’s built-in command FindGeometricTransform was used to transform detected spot coordinates. The correct transformation function was determined by aligning the positions of 100 nm diameter Tetraspeck beads (fluorescent in all channels) imaged on the same day of each experiment. The transformation function was only used to correct for spot positions so that spots could be properly categorized, as described above. It was not used to correct the raw images themselves, as this could lead to artifactual image distortions when reassigning pixel values. Therefore, a slight offset between channels can be observed in the raw and 2D max-projected images and movies.

To calibrate the intensity of single-mRNA translation signals to the number of mature proteins (a proxy for the number of ribosomes translating the mRNA), the average fluorescence signals of the spaghetti-monster-tagged firefly luciferase nonoptimal and optimal reporters were compared to that of spaghetti monster-β-Actin-MS2, a reporter that was previously shown to have an average fluorescence equivalent to 11.4 ± 2.0 SM-tagged proteins (or ribosomes) in U2OS cells (Koch et al., 2020). To make the comparison as fair as possible, cells were imaged in two chambers on the same day with the same imaging conditions (50 mW for 561 nm and 15 mW for 637 nm laser). In the first chamber, cells were bead loaded with α-Flag Fab (Cy3) and spaghetti monster- β-Actin-MS2. In the second chamber, cells were bead loaded with α-Flag Fab (Cy3) and either of the spaghetti-monster-tagged firefly luciferase optimality reporters.

#### Ribo-seq library preparation

Libraries were prepared according to Calviello et al., 2016. Briefly, cells were coated with ice-cold PBS supplemented with 100 μg/mL cycloheximide, before immediately aspirating wash and dipping the bottom of each plate in LN_2_. Ice-cold lysis buffer B (LyB: 20 mM Tris HCL, pH 7.4, 150 mM NaCl, 5 mM MgCl2, with freshly added 1 mM DTT, 100 µg/mL cycloheximide [Sigma], 1% (v/v) Triton X-100, and 25U/mL TURBO DNase [Invitrogen]) was dripped onto frozen dishes. Plates were thawed on ice, and the lysate was collected by scrapping and transferred to a pre-cooled 1.5 tube. Lysates were incubated on ice for ten minutes, triturated ten times with a 26-gauge needle, and centrifuged ten minutes, 20,000 x g at 4°C. Clarified lysate was transferred to pre-cooled tubes, flash frozen in LN_2_, and stored at –80°C. Ribo-seq experiments were performed in biological triplicate.

RNA lysate concentration was measured using the Qubit RNA BR kit. 85 μg lysate was footprinted for ribosomal fragments with 750 U RNase I (Ambion) and brought to 300 µL with 1x lysis buffer B. Samples were incubated for 45 min, shaking at 300 rpm at room temperature. Ten µL SUPERase•In (Ambion) was added to stop digestion, and samples were placed on ice. MicroSpin S-400 HR columns (GE Healthcare) were equilibrated by removing storage buffer and washing with PB buffer (20 mM Tris HCl, pH 7.4, 150 mM NaCl, 5 mM MgCl2, and 1 mM DTT and 100 µg/mL cycloheximide [Sigma]). 100 µL digested lysate was loaded per spin, and flowthrough was transferred to a pre-chilled five mL tube. Three volumes of TRIzol LS were added before purification using Direct-zol RNA Micro-Prep Kit (with in-column DNase treatment), and RNA was eluted with 11 µL RNase-free water. Five µg of RNA was ribo-depleted using siTOOLs Human RiboPOOL for Ribosome Profiling (Galen Molecular), following manufacturer’s instructions. Ribo-depleted RNA was precipitated in ethanol overnight at -80°C and resuspended in RNase-free water. Next, 27-30 nt fragments were recovered from 15% TBE-Urea PAGE gel, using 27 nt and 30 nt ribomarkers as guides (Ribo27: 5’ rArUrG rUrArC rArCrG rGrArG rUrCrG rArGrC rUrCrA rArCrC rCrGrC/3Phos/; Ribo30: 5’ rArUrG rUrArC rArCrG rGrArG rUrCrG rArGrC rUrCrA rArCrC rCrGrC rArArC/3Phos/). Bands were visualized using 1x SybrGold (Invitrogen). Extracted gel was shredded by centrifugation and incubated in RNA extraction buffer (400 mM NaCl, 1 mM EDTA, and 0.25% (wt/v) SDS) at room temperature, rotating, for 2.5 hours, before being filtered through 0.45 μm cellulose acetate Spin-X filter (Corning) at top speed. RNA was precipitated with isopropanol overnight at –80°C and resuspended in RNase-free water. Samples were treated with PNK, precipitated with isopropanol overnight at –80°C, and resuspended in RNase-free water.

Libraries were prepared from fragments using QIASeq miRNA Library preparation kit (Qiagen), with indexes supplied in QIAseq miRNA NGS 96 Index IL plate kit (Qiagen). Pilot PCRs using HotStarTaq 2X Mastermix (Qiagen) were performed and run out on 6% TBE-PAGE gels with 186bp Qiagen marker PCR product to best determine cycle number. After amplification, samples were purified according to manufacturer instructions, and libraries were eluted in RNase-free water. The quality of the libraries was determined with Qubit dsDNA HS assay and Tapestation D1000 HS Screentape (Agilent). Ribo-seq libraries were sequenced on NovaSeq6000 Illumina sequencer at the Shared Genomics Core at University of Colorado Anschutz.

#### RNA-seq

10% of each lysate from the Ribo-seq sample was stored at -80°C with three volumes of TRIzol LS. Purified RNA input sample was ribo-depleted using hybridizing rRNA complementary DNA oligo mix (final conc 16.5 µM) and thermostable RNaseH (10 U), as previously described (Baldwin et al., 2021). Samples were purified with 2.2X RNA Clean XP beads and eluted with RNase-free water. Cleaned samples were DNase-treated with TURBO DNase (Invitrogen), incubating at 37°C for thirty minutes. The samples were cleaned again using 2.2X RNA Clean XP beads and eluted in 22 µL 1x Elute Prime Fragment Buffer included in KAPA RNA Hyperprep library kit. Samples were fragmented by heating at 85°C for six minutes. RNA-seq libraries were generated per KAPA RNA Hyperprep library kit protocol, with Illumina compatible IDT-UBI adapters (KAPA Dual-Indexed Adapter Kit [Roche]) cDNA was amplified for ten cycles. PCR products were purified with 0.8X bead-based cleanup (KAPA Pure beads) and eluted in 10 mM Tris-HCl (pH 8.0 – 8.5). Samples were checked for quality control with Qubit dsDNA HS kit (Invitrogen) and Tapestation HS D1000 Screentape (Agilent). Libraries were sequenced on NovaSeq6000 Illumina sequencer at the Shared Genomics Core at University of Colorado Anschutz.

#### Ribo-seq data processing

UMI-containing reads (Read 2 only) were extracted from the fastq files using UMI-tools (Smith et al., 2017). Next, adapters were trimmed, and reads shorter than 18 nts were filtered with Cutadapt (Martin, 2011). The trimmed fastq files were reverse complemented, and any remaining rRNA-mapped reads were filtered out using Bowtie (version 1.2.2) and SAMtools (version 1.12) (Langmead et al., 2009, Danecek et al., 2021. The remaining reads were aligned and mapped to the human genome (hg38) with STAR (version 2.6.0a) (Dobin et al., 2013). The output .bam files were de-duplicated using UMI-tools and transcriptome-wide transcript expression was calculated with Salmon (version salmon 1.6.0) (Patro et al., 2017). The transcriptome used was Gencode v26. Bigwig files were also generated using bamCoverage to visualize RNA-seq and Ribo-seq coverage across the genome. The libraries were further checked for quality control with ribosomeProfilingQC R package (Ou and Hoye, 2022), where the following were analyzed: 1) average read length (27-30 nts for ribosome-associated fragments expected), 2) percentage of reads mapping to CDS, 3) the mean of coverage across the CDS, and 4) proper phasing across the ORFs (Supplemental file 1). Phasing plots were generated with Ribo-seq fragments of 27, 28, and 29 nts in length.

To analyze the translational efficiency of our reporters, we extracted the reads that did not map to the genome from the output .bam files. These reads were de-duplicated using UMI-tools (ribo-seq only) and mapped to the reporter sequences using Bowtie (version 1.2.2). Each condition included the fasta sequence from the transfected 1) *Renilla* luciferase plasmid and 2) optimal or nonoptimal dual-tagged firefly luciferase reporter plasmids. Bedtools coverage (version v2.26.0- 129-gc8b58bc; Quinlan and Hall, 2010) and dplyr tools in R were used to quantify reads across each reporter. RPKM was measured using the equation “RPKM = numReads / (geneLength/1000 * totalNumReads/1,000,000)” in R and Excel. The P-site was estimated to be 13 nucleotides downstream of the 5’-end of each read. Additional analysis was performed using in-house scripts in R.

### QUANTIFICATION AND STATISTICAL ANALYSIS

An indicated in figure legends and text, built-in R studio Student’s t-test or Mann-Whitney U tests were used to calculate statistical significance. Unless otherwise stated, n refers to the number of biological replicates.

**KEY RESOURCES TABLE** – see attached document

**Movie S1. Co-labeled HaloTag-MCP and anti-FLAG Cy3 signal on the nonoptimal firefly luciferase reporter shows sites of active translation.** To confirm that co-labeled signal were sites of active translation on the nonoptimal firefly luciferase reporter, cells were treated with 50 ug/mL puromycin, and anti-Flag Cy3 signal can be seen rapidly dissociating from each mRNA puncta. The field of view is 512 x 512 pixels with 130 nm per pixel. Image settings are further described in the methods. The movie is a max-Z projection with one frame every six seconds, and puromycin was added at frame 5.

**Movie S2. Co-labeled HaloTag-MCP and anti-FLAG Cy3 signal on the optimal firefly luciferase reporter shows sites of active translation.** To confirm that co-labeled signal were sites of active translation on the optimal firefly luciferase reporter, cells were treated with 50 ug/mL puromycin, and anti-Flag Cy3 signal can be seen rapidly dissociating from each mRNA puncta. The field of view is 512 x 512 pixels with 130 nm per pixel. Image settings are further described in the methods. The movie is a max-Z projection with one frame every six seconds, and puromycin was added at frame 5.

**Movie S3. Representative movie illustrating nascent chain tracking with the nonoptimal firefly luciferase reporter.** Movie showing spots of active translation (co-labeled HaloTag-MCP and anti-Flag Cy3 signal) on the nonoptimal firefly luciferase reporter. The field of view is 512 x 512 pixels with 130 nm per pixel. Image settings are further described in the methods. The movie is a max-Z projection with one frame every six seconds.

**Movie S4. Representative movie illustrating nascent chain tracking with the optimal firefly luciferase reporter.** Movie showing spots of active translation (co-labeled HaloTag-MCP and anti-Flag Cy3 signal) on the optimal firefly luciferase reporter. The field of view is 512 x 512 pixels with 130 nm per pixel. Image settings are further described in the methods. The movie is a max-Z projection with one frame every six seconds.

**Table S1. Expanded list of RT-qPCR primers used, as referred in STAR Methods Key Resource Table.**

---

**Figure S1.**
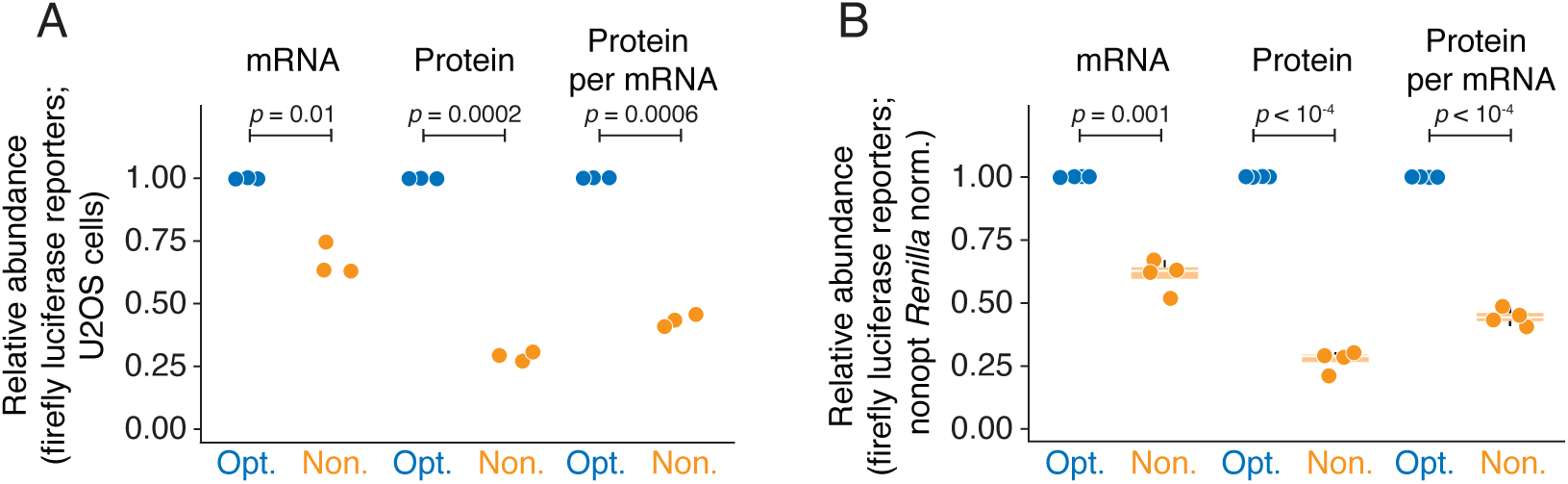
Codon optimality also has protein-level effect in U2OS cells and is independent of transfection control. (A) In U2OS cells, firefly luciferase optimality reporters show differences in protein output, even whennormalized for transcript abundance. U2OS cells were transfected with either Flag-tag spaghetti-monster optimal (Opt.) or nonoptimal (Non.) firefly luciferase reporters and *Renilla* luciferase, as a transfection control. mRNA levels were quantified by qRT-PCR, and protein activity was quantified by dual luciferase assay. Shown are scatter plots for the fold-change of the nonoptimal reporter relative to the optimal one for mRNA abundance, protein abundance, and normalized protein-per-mRNA abundance. P-values were determined using a paired Student’s t-test. (B) Additional protein level effect does not depend on co-transfected optimized *Renilla* luciferase. HEK293T cells were transfected with either Flag-tag spaghetti-monster optimal (Opt.) or nonoptimal (Non.) firefly luciferase reporters and nonoptimal *Renilla* luciferase, as a transfection control. Otherwise,as in Figure S1A.

**Figure S2.**
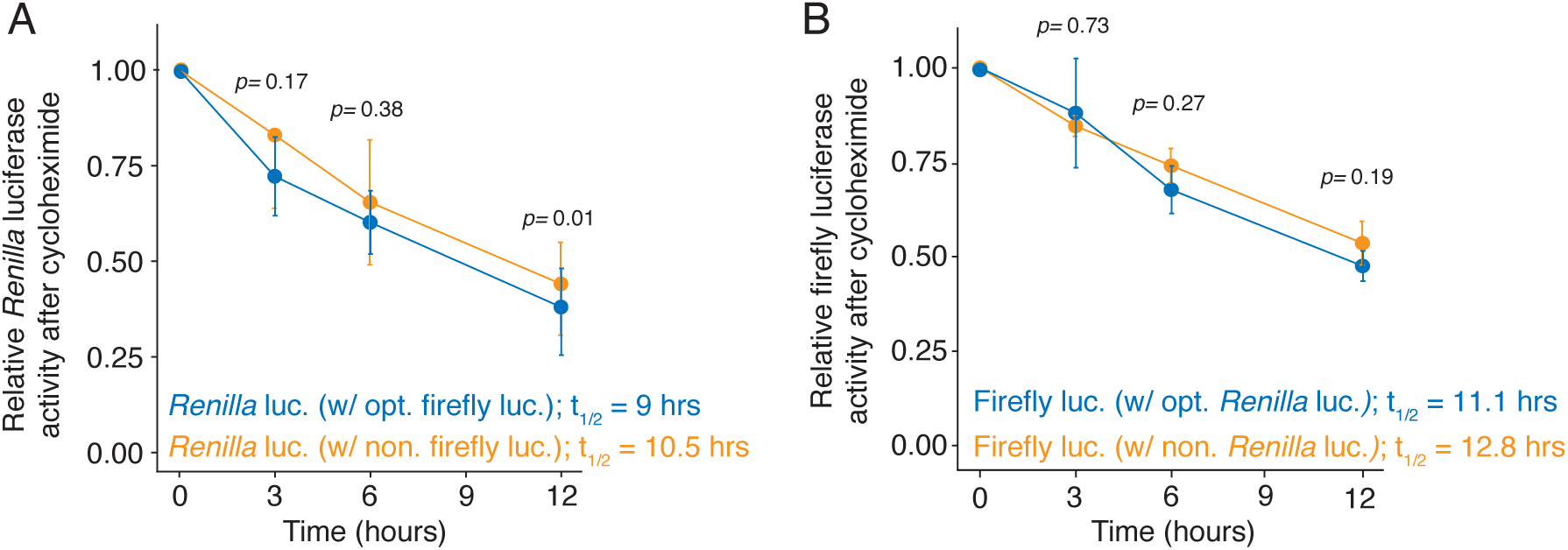
Cycloheximide time course shows no difference in the half-lives of either co-transfection control. (A) Optimized *Renilla* luciferase co-transfection controls from Figure 2B have no significant difference in half-life. Cells were co-transfected with the spaghetti-monster firefly luciferase reporters and optimized *Renilla* luciferase, and then harvested at various time points following cycloheximide treatment. The amount of *Renilla* luciferase control was determined by luciferase assay and normalized to the initial (0 hr.) time point. P-values were determined using a paired Student’s t-test. (B) Optimized firefly luciferase co-transfection controls from Figure 2C have no significant difference in half-life. Cells were co-transfected with the spaghetti-monster *Renilla* luciferase reporters and optimized firefly luciferase, and then harvested at various time points following cycloheximide treatment. The amount of firefly luciferase control was determined by luciferase assay and normalized to the initial (0 hr.) time point. P-values were determined using a paired Student’s t-test.

**Figure S3.**
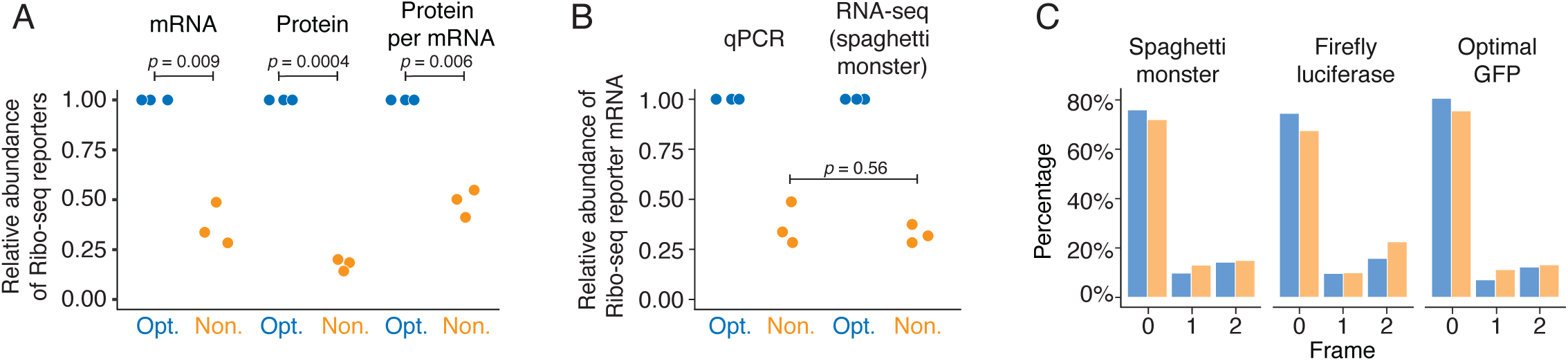
Ribo-seq reporters show additional protein-level effect, for which frameshifting is not responsible. (A) Dual-tagged Ribo-seq optimality reporters show differences in protein output, even after controlling for transcript levels. HEK293T cells were transfected with either nonoptimal (Non.) or optimal (Opt.) firefly luciferase, tagged at the N-terminus with the spaghetti-monster tag, and tagged with the C-terminus with GFP. Transcript abundance was quantified by qRT-PCR, and protein activity was quantified by dual luciferase assay. Shown are scatter plots for the fold-change of the nonoptimal reporter relative to the optimal one for mRNA abundance, protein abundance, and normalized protein-per-mRNA abundance. P-values were determined using a paired Student’s t-test. (B) Nonoptimal codons reduce the mRNA abundance of the dual-tagged reporters, as determined by spaghetti-monster mapped reads. mRNA abundance for each reporter was determined by RT-qPCR or RNA-seq (from the same samples used for the Ribo-seq libraries) and normalized to the Renilla luciferase co-transfection control. Shown are scatter plots for the fold-change of the nonoptimal reporter relative to the optimal one. RNA-seq reads quantified were mapped to spaghetti-monster (SM) tag only. P-values were determined by unpaired Student’s t-tests. (C) There is no evidence of substantial frameshifting due to the nonoptimal coding region. Shown is the frame (zero, one, and two) of Ribo-seq reads in each section of the dual-tagged reporter: N-terminal spaghetti-monster, optimal or nonoptimal firefly luciferase, and C-terminal GFP. Blue color refers to the optimal firefly luciferase reporter, while orange refers to the nonoptimal one. The first 25 codons of the spaghetti-monster tag and the last 25 codons of the GFP tags were excluded from the analysis. The three Ribo-seq replicates were pooled.

**Figure S4.**
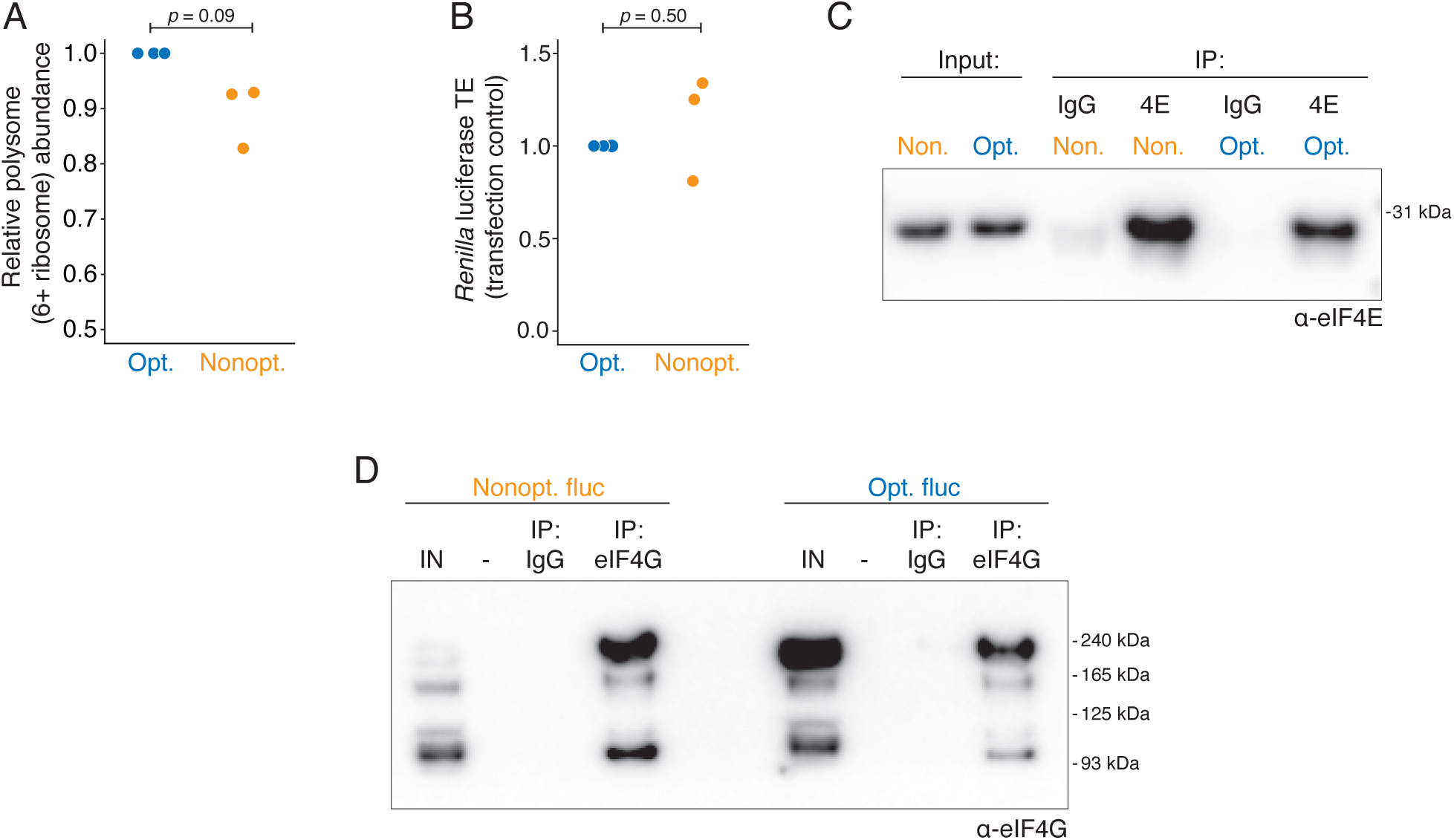
There is reduced initiation on nonoptimal transcripts. (A) Nonoptimal codon usage decreases association with the polysome fraction. Plotted is the abundance of the nonoptimal firefly reporter in the polysome fraction (six or more ribosomes) relative to the optimal reporter. The p-value was determined using a paired Student’s t-test. (B) The translational efficiency of co-transfected*Renilla* luciferase with dual-tagged reporters was not significantly different between the conditions. Translational efficiency was defined as Ribo-seq reads relative to RNA-seq abundance. *Renilla* luciferase co-transfected with optimal dual-tagged firefly luciferase reporter was normalized to 1. P-value was determined by paired Student’s t-test. (C) eIF4E was immunoprecipitated equally between both samples. Western blotting was performed, confirming successful eIF4E immunoprecipitation for both conditions: nonoptimal firefly luciferase (Non.), and optimal firefly luciferase (Opt.). Estimated size of eIF4E is 28 kDa. (D) As in B, except for eIF4G RNA immunoprecipitations. Estimated size of eIF4G is 220 kDa.

